# Constitutive androstane receptor directs developing Th1 cells towards the Tr1 lineage

**DOI:** 10.64898/2026.07.23.740345

**Authors:** Courtney L. Hegner, Blake M. Frey, Akshaya Balasubramanian, Henry D. Dionne, Huijuan Yang, B. JoNell Hamilton, Casey T. Weaver, Mark S. Sundrud

## Abstract

Constitutive androstane receptor (CAR; encoded by *Nr1i3*) is a nuclear xenobiotic receptor that mediates hepatic drug and bile acid metabolism. We previously identified CAR as also operating in CD4^+^ T helper (T_H_) cells, where CAR-dependent gene expression mitigates bile acid toxicity and promotes a Foxp3^-^IL-10^+^type 1 regulatory (Tr1)-like phenotype in the small intestine. Here, we show that CAR acts early and specifically during the priming of type 1 immune responses to stabilize Tr1 lineage commitment. CAR-dependent Tr1 cells formed during type 1 (Th1-associated) but not type 3 (Th17-associated) intestinal inflammation. *In vitro*, IL-27 upregulated CAR expression during naïve T_H_ cell activation, which contributed to *Il10* induction. Single-cell analysis revealed that naïve T_H_ cells primed with IL-27 adopt a multipotent “Th1/Tr1 precursor” (THR1p) transcriptional state, which subsequently diverges into Th1 or Tr1 developmental trajectories. CAR transcriptional activity peaked in THR1p cells, upregulating Tr1 genes, including *Il10*, and repressing Th1 genes. Moreover, glucocorticoid receptor activation—which increases CAR expression in hepatocytes—synergized with IL-27 to augment both CAR expression and CAR-dependent Tr1 differentiation. Together, these results suggest that CAR acts in a lineage-biased manner to enforce Tr1-mediated immune tolerance during type 1 intestinal inflammation, and this pathway is amplified by glucocorticoids.

## Introduction

Constitutive androstane receptor (CAR; encoded by *Nr1i3*) is a member of the nuclear receptor (NR) super-family known for promoting phase I/II metabolism of potentially toxic compounds in the liver, including synthetic drugs and bile acids^1,2^. Unlike most NRs, the CAR ligand-binding domain (LBD) contains an extended AF-2 motif that confers high basal transcriptional activity in the absence of bound ligand. Thus, CAR regulation in hepatocytes involves phosphorylation-dependent cytoplasmic sequestration; compounds known to activate CAR (*e.g*., phenobarbital, bile acids, etc.) do so indirectly by promoting CAR dephosphorylation and nuclear translocation^3,4^. Once in the nucleus, CAR binds xenobiotic response elements (XREs)—together with RXRα or as a monomer— and promotes transcription of target genes, which in hepatocytes include cytochrome P450 (CYP) enzymes and efflux transporters that coordinate the modification and biliary excretion of toxic small molecules^1,2^.

We previously discovered that CAR also regulates functions of CD4^+^ T helper (T_H_) cells in the lamina propria of the distal small intestine (*i.e.*, ileum)^5^. As the sole and dedicated site of intestinal bile acid absorption, the ileal mucosa is a unique intestinal microenvironment where resident or infiltrating immune cells are exposed to potentially toxic bile acid concentrations^5^. Our results supported a model in which CAR activates complementary transcriptional programs within ileal T_H_ cells, including: (*i*) a ‘hepatocyte-like’ program, including detoxifying enzymes (*e.g*., *Cyp2b10*) and efflux transporters (*e.g*., *Abcb1a*), which support T_H_ cell homeostasis in the presence of bile acids; and (*ii*) a type 1 regulatory (Tr1)-like program, including *Il10*, that enforces ileal immune tolerance^5^.

Tr1 cells are an important subset of CD4^+^ T cells that express IL-10, but not FOXP3^6,7^. Tr1 cells were first identified in humans with severe combined immunodeficiency (SCID) that displayed long-term immune tolerance toward HLA-mismatched hematopoietic stem cell transplantation (HSCT)^6,8^. Subsequent studies have shown that *bona fide* Tr1 cells: (*i*) are found in humans and mice; (*ii*) are distinguished from other IL-10-secreting T cells by the co-expression of multiple co-inhibitory receptors (*e.g*., CD49b, LAG3, PD-1, CTLA-4); and (*iii*) display both IL-10- and contact-mediated immune suppression *in vitro* and *in vivo*^7,9,10^. Tr1 cells have been observed in many tissues—including small intestine lamina propria—but arise mostly in settings of chronic infection, cancer and autoimmunity^7,9-14^. It remains controversial if Tr1 cells reflect a single developmental lineage, or a heterogenous collection of functionally and phenotypically similar cells with distinct developmental histories. On one hand, *bona fide* Tr1 cells can develop directly from antigen-stimulated naïve T_H_ cells in the presence of IL-10^15^, or the IL-12 family cytokine, IL-27^16-18^. On the other hand, more recent studies suggest that Tr1-like cells may also *trans*-differentiate from other effector (T_EFF_) subsets, including Th1, T follicular helper (T_FH_) and Th17 cells^7,14,19-21^. Tr1 cells lack a single lineage-defining transcription factor, although networks involving BATF, IRF1, STAT3, c-Maf, Blimp-1 and AhR are consistently implicated in Tr1 cell development and *Il10* locus control^7,22^.

Here, we show that CAR acts earlier and more specifically than anticipated during the priming of type 1 (Th1-associated) intestinal inflammatory responses to steer developing Th1 cells towards the Tr1 lineage. CAR expression was induced during naïve T_H_ cell activation by IL-27 signaling and supported IL-27-induced *Il10* upregulation. Both *in vivo* during intestinal type 1 inflammation and *in vitro* during naive T_H_ cell activation in the presence of IL-27, CAR transcriptional activity peaked within a nascent activated T_H_ cell subset with multipotent potential, which we term Th1/Tr1 precursor (THR1p), and which gives rise to mature intestinal Th1 and Tr1 cells. Within THR1p, CAR amplified Tr1-associated genes, including *Il10*, and downregulated Th1-associated genes. Furthermore, activation of glucocorticoid receptor (GR)—known to *trans*-activate CAR expression in hepatocytes—synergized with IL-27 signaling to augment Tr1 cell development in a CAR-dependent manner. Thus, rather than governing tolerogenic functions of mature Tr1 cells locally in the ileum, our results suggest an alternative model in which CAR acts as a molecular rheostat in secondary lymphoid organs to dictate the balance between Th1-mediated immunity and Tr1-mediated tolerance. CAR-dependent Tr1 cells may be a dedicated regulatory counterpart to effector Th1 cells—akin to the reciprocal relationship between FOXP3^+^ Treg and effector Th17 cells—and endogenous glucocorticoids (produced in response to intestinal inflammation^23,24^) may provide an orthogonal leverage-point to augment CAR-dependent Tr1 cell development.

## Results and Discussion

### CAR promotes intestinal Tr1 cell development during type 1 inflammatory responses

We have shown that CAR steers development of small intestinal Foxp3^-^IL-10^+^ Tr1-like cells in the naive T cell transfer mouse colitis model^5^. Small intestinal Tr1-like cells in this model display CAR-dependent expression of genes previously ascribed to IL-27-induced Tr1 cells *in vitro*, and accumulation of CAR-dependent Tr1-like cells in the ileum suppresses bile acid-induced intestinal inflammation^5,7,14,18^. To understand the broader inflammatory features that associate with CAR-dependent Tr1 cell development during T cell transfer colitis, we performed *ex vivo* flow cytometry on wild type or CAR-deficient (*Nr1i3*^-/-^) T_H_ cells recovered from ileums of *Rag1*^-/-^ recipients 2-weeks post-mixed naïve T cell transfer. To streamline detection of IL-10-expressing donor T_H_ cells, we crossed CD45.1/CD45.2-mismatched wild type or CAR-deficient donor mice with *Il10*-IRES-Thy1.1 reporter (*i.e*., 10BiT) mice^25^. Consistent with our prior results, a sizeable portion of wild type T_H_ cells recovered from ileums of transplanted *Rag1*^-/-^ mice adopted a Foxp3^-^I0BiT^+^ Tr1-like phenotype; these cells were also evident in colons and spleens of the same animals, albeit at lower frequencies. (**Fig. 1A-B**). Ileal Foxp3^-^I0BiT^+^ Tr1-like cells also produced IL-10 upon *ex vivo* stimulation and showed strict CAR-dependent expression of both the 10BiT reporter and endogenous IL-10 (**Fig. 1A-B**, **S1A-D**). Virtually all Tr1-like and non-Tr1 cells in this model expressed the canonical Th1-associated cytokine, IFNγ, whereas few cells expressed Th17 (IL-17A) or Th2 (IL-4) cytokines (**Fig. 1A-B, S1E-G**, data not shown). CITE-seq (Cellular Indexing of Transcriptomes and Epitopes by Sequencing) analysis of *ex vivo*-isolated ileal T_H_ cells confirmed that Foxp3^-^I0BiT^+^ cells (cluster 3 in **Fig. 1C-D**) expressed prototypical Tr1-associated co-inhibitory receptors [*Ctla4*, *Pdcd1* (PD-1), *Havcr2* (Tim3), *Lag3*], granzymes (*Gzmk*) and transcription factors [*Stat3*, *Maf*, *Prdm1*, *Tox*, *Eomes*]^7,9,10,14,20,22,26,27^.

**Figure 1:**
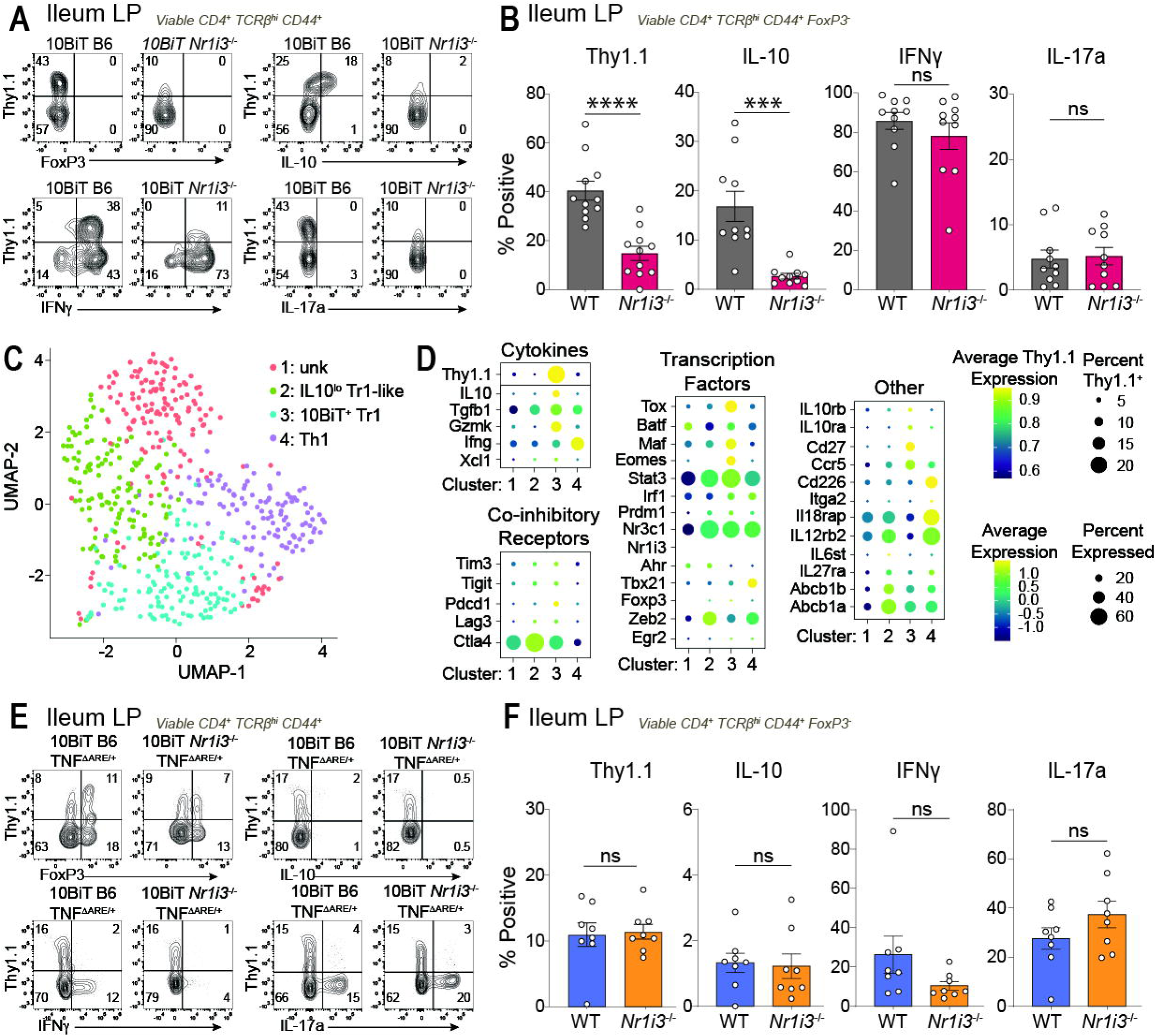
CAR promotes intestinal Tr1 cell development during type 1 inflammatory responses. **(A-B)** *in vivo* transfer of CAR-sufficient (WT) and CAR-deficient (*Nr1i3^-/-^*) 10BiT naïve CD4^+^ T cells into Rag1-deficient (Rag1^-/-^) hosts. Cells were isolated from the ileum 2.5 weeks post transfer, stained and analyzed via flow cytometry. Cells were gated on CD4^+^ T_H_ cells and gated on congenic marker, CD45.1^+^ WT and CD45.1^-^ *Nr1i3^-/-^*. **(A)** Representative flow cytometry plot of Ileal lamina propria (LP) T_H_ cells gated on viable CD4^+^ TCRβ^hi^ CD44^+^; FoxP3, IL-10, IFNγ, or IL-17a vs Thy1.1. **(B)** Ileal expression of Thy1.1 10BiT, IL-10, IFNγ, and IL-17a gated on CD44^+^ FoxP3^-^ was measured. (Percent; n=11; ± s.e.m.) \*\*\**P<*0.001, \*\*\*\**P*<0.0001, paired, parametric two-tailed Student’s *t-*test. **(C-D)** CITE-seq analysis of ileal CD4^+^ T_H_ cells isolated post adoptive transfer in Rag1^-/-^. **(C)** UMAP projection of 4 ileal subclusters. **(D)** Expression of key Tr1- and Th1-associated genes in 4 ileal subclusters. **(E-F)** TNF^ΔARE/+^ mice crossed to 10BiT CAR-sufficient and CAR-deficient mice were cohoused until 20 weeks of age. Cells were isolated from the ileum, stained and analyzed via flow cytometry. **(E)** Representative flow cytometry plot of Ileal lamina propria (LP) T_H_ cells gated on viable CD4^+^ TCRβ^hi^ CD44^+^; FoxP3, IL-10, IFNγ, or IL-17a vs Thy1.1. **(F)** Ileal expression of Thy1.1 10BiT, IL-10, IFNγ, and IL-17a gated on CD44^+^ FoxP3^-^ was measured. (Percent; n=8; +s.e.m.), paired, one-way Anova with Greisser-Greenhouse correction.

To determine if CAR promotes ileal Tr1 cell development in other inflammatory contexts, we analyzed CAR-dependent ileal Tr1 cell development in *Tnf*^ΔARE/+^ mice, which spontaneously develop Crohn’s disease-like ileitis due to ablation of a destabilizing 69-bp A/U-rich element in the 3’UTR of the endogenous *Tnf* locus^28^. *Tnf*^ΔARE/+^ mice used here were crossed with CAR-sufficient or CAR-deficient 10BiT reporter animals; ileal Tr1 cells were analyzed in control or CAR-deficient littermates that were cohoused at weaning—to normalize microflora exposure—and maintained until 20-wks of age, when 100% of *Tnf*^ΔARE/+^ mice in our colony displayed moderate-to-severe ileitis. Indeed, Foxp3^-^I0BiT^+^ Tr1-like cells were also evident in ileums of *Tnf*^ΔARE/+^ mice, whereas all 10BiT^+^ T_H_ cells found in spleens and colons of these mice were Foxp3^+^ Tregs (**Fig. 1E, S1H-K)**. Unlike the T cell transfer colitis model, however, loss of CAR had no bearing on the number or frequency of ileal Tr1-like cells in *Tnf*^ΔARE/+^ mice, and both Tr1-like (10BiT^+^) and bystander non-Tr1 (10BiT^-^) cells expressed more IL-17A than IFNγ (**Fig. 1E-F, S1L-N**). These results suggest that CAR-dependent Tr1 cell development occurs principally in settings of type 1 intestinal inflammation.

### CAR-dependent Tr1 cells and inflammatory Th1 cells derive from a common activated progenitor

To define the development, heterogeneity, and CAR-dependency of ileal Tr1 cells during T cell transfer colitis, we expanded our CITE-seq analysis. We noted that *bona fide* I0BiT^+^ Tr1 (cluster 3) cells were present in the ileum together with a second cluster of *Il10*^lo^ Tr1-like (cluster 2) cells, as well as with inflammatory Th1 (cluster 4) cells (**Fig. 1C-D, 2A**). A fourth cluster of activated cells (cluster 1) remained unannotated after gene set enrichment analysis (GSEA) and examination of known marker genes (**Fig. 1C-D**). *Il10*^lo^ Tr1-like (cluster 2) cells were 10BiT^-^ and expressed low levels of Tr1-related co-inhibitory receptors and transcription factors but showed enrichment for gene expression previously associated with IFNγ^+^IL-10^+^ Tr1 cells from mice infected with malaria parasites (**Fig. 1C-D, 2A**)^29^. Conversely, *bona fide* I0BiT^+^ Tr1 (cluster 3) cells showed enrichment of genes previously found in Eomes^+^ Tr1 cells from mice with chronic lymphocytic leukemia (CLL)-like disease (**Fig. 2A**)^14^. Eomes is a T-box family transcription factor (TF) related to the Th1 master TF, T-bet, and was first identified for promoting cytotoxic functions in CD8^+^ T and natural killer (NK) cells^30,31^. However, more recent studies have implicated Eomes in the regulation of both Th1 and Tr1 cells^7,14,26,27^. These results reveal a range of developmentally related Th1 and Tr1 cell fates generated during type 1 intestinal inflammatory responses.

**Figure 2:**
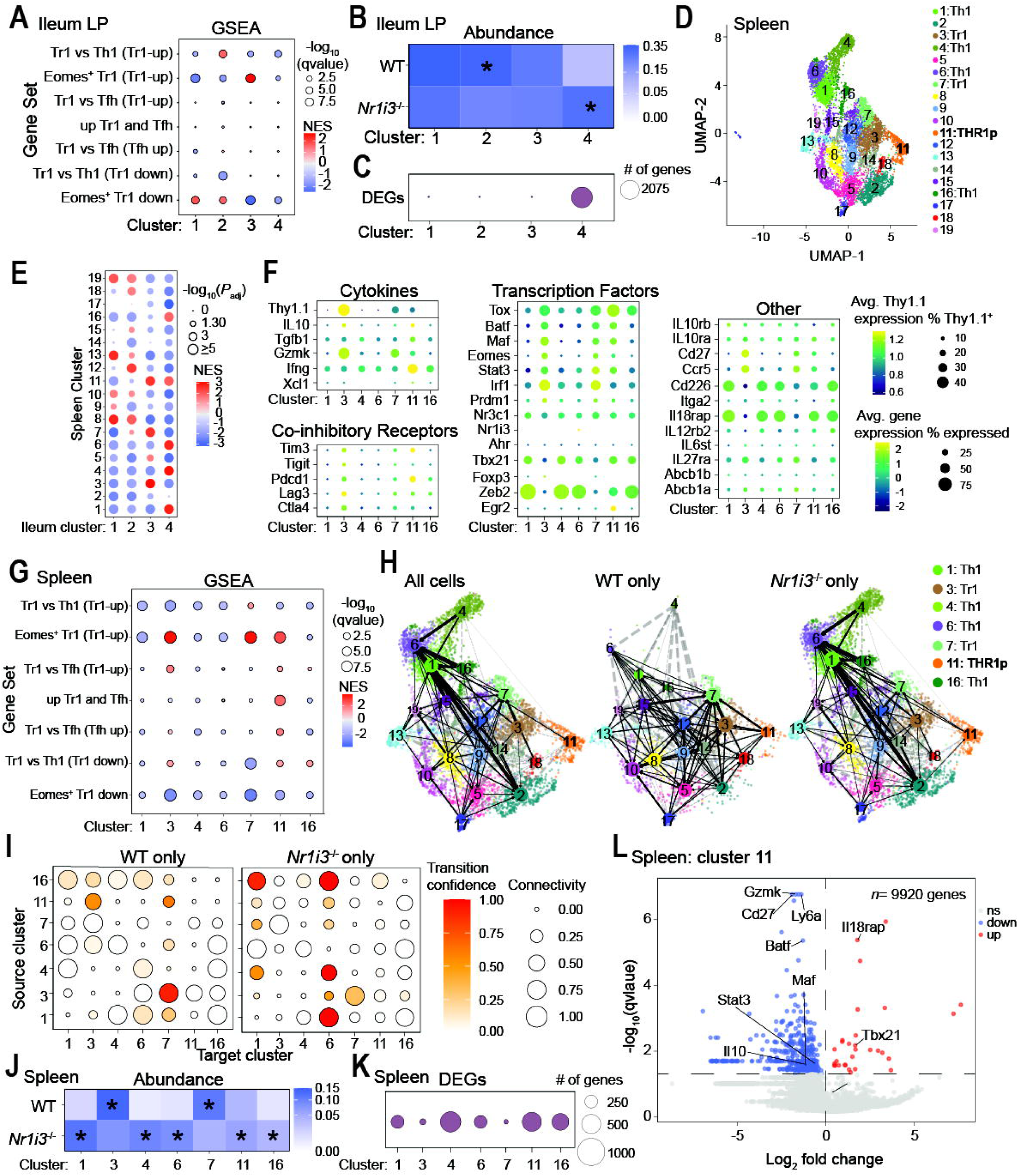
CAR-dependent Tr1 cells derive from a common progenitor with inflammatory Th1 cells. **(A-C)** CITE-seq analysis of ileal CD4^+^ T_H_ cells isolated post adoptive transfer in Rag1^-/-^. **(A)** Geneset enrichment analysis (GSEA) of ileal clusters against published Tr1 genesets. **(B)** Abundance of ileal WT (10BiT. B6) and CARko (*Nr1i3^-/-^*) cells in each cluster. **(C)** Differentially expressed genes (DEG) between *Nr1i3^-/-^* and WT in each ileal cluster. **(D-I)** CITE-seq analysis of splenic CD4^+^ T_H_ cells isolated post adoptive Rag1^-/-^ transfer. **(D)** UMAP projection of 19 spleen subclusters. **(E)** GSEA between ileal and spleen subclusters. **(F)** Expression of key Tr1- and Th1-associated genes in Th1, Tr1, and THR1p subclusters. **(G)** Geneset enrichment analysis (GSEA) of spleen Th1, Tr1, and THR1p subclusters against published Tr1 genesets. **(H-I)** Partition-based graph abstraction (PAGA) velocities of all splenic sublusters with total, just WT, or just *Nr1i3^-/-^* cells. **(H)** PAGA velocity UMAPs **(I)** PAGA connectivies and transition confidence in Th1, Tr1, and THR1p subclusters. **(J)** Abundance of ileal WT (10BiT. B6) and CARko (*Nr1i3^-/-^*) cells in Th1, Tr1, and THR1p subclusters. **(K)** Differentially expressed genes (DEG) between *Nr1i3^-/-^*and WT in Th1, Tr1, and THR1p subclusters. **(L)** DEG in THR1p (spleen cluster 11) between *Nr1i3^-/-^* and WT.

Loss of CAR altered the proportions of Tr1 and Th1 cells in the ileum but had little impact on gene expression within these differentiated subsets. For example, nearly 60% of all CAR-sufficient ileal T_H_ cells fell within the two Tr1 cell clusters (cluster 2, 3), whereas this dropped to ∼40% among CAR-deficient T_H_ cells (**Fig. 2B**). Conversely, only ∼7% of CAR-sufficient ileal T_H_ cells were Th1 (cluster 4) cells, but this rose to almost 30% for T_H_ cells lacking CAR (**Fig. 2B**). Finally, CAR/*Nr1i3* expression was not detectable in any ileal Tr1 or Th1 cell cluster, and few differentially expressed genes were observed between CAR-sufficient and CAR-deficient ileal Tr1 clusters (**Fig. 1D, 2C**). Thus, we postulated that CAR might operate at earlier stages of T_H_ cell lineage commitment to direct the development, rather than function, of intestinal Tr1 cells.

To test this hypothesis, we expanded our CITE-seq studies to analyze transcriptional behaviors of CAR-sufficient and CAR-deficient 10BiT reporter T_H_ cells from spleens of the same transplanted *Rag1*^-/-^ recipient mice, assuming that cells in spleen include those primed by antigen in either mesenteric lymph nodes or Peyer’s patches and returning to the intestinal lamina propria via peripheral circulation. Sub-clustering of all CAR-sufficient and CAR-deficient spleen T_H_ cells distinguished 19 clusters, among which clusters 1/4/6/16 and 3/7, respectively, displayed transcriptional characteristics of ileal Th1 and Tr1 cells (**Fig. 2D-G**). More interestingly, a single cluster of spleen cells (cluster 11) displayed enrichment for both Tr1- and Th1-associated gene expression (**Fig. 2D-G**). Partition-based graph abstraction (PAGA) of RNA velocities for all spleen T_H_ cell clusters revealed that cluster 11 was a branch point from which Tr1 and Th1 developmental trajectories diverged (**Fig. 2H-I**). Similar to cells found in the ileum, loss of CAR altered the abundance of Tr1 and Th1 cells in the spleen, whereas a more equal ratio of WT and CAR-deficient cells was found in the shared Th1/Tr1 cluster (cluster 11) (**Fig. 2J**). Importantly, cluster 11 cells displayed a high number of differentially expressed genes, when comparing CAR-sufficient and CAR-deficient constituents (**Fig. 2K**), suggesting that CAR is transcriptional active this uniquely multi-potent developmental state. Specifically, cluster 11 cells lacking CAR displayed reduced expression of *Il10*, as well as of key Tr1-associated TFs (*e.g*., *Stat3*, *Batf*, *Maf*, *Irf1*, *Eomes*) and co-inhibitor receptors (*e.g*., *Lag3*, *Pdcd1*), but expressed higher levels of Th1-assocaited transcripts, including *Tbx21* (T-bet) and *Il18rap* (**Fig. 2L**). Accordingly, cluster 11 cells differentiated along a Tr1 trajectory (*i.e*., towards clusters 3, 7) in the presence of CAR, whereas these cells diverted towards the Th1 lineage (*i.e*., towards clusters 1, 6) in the absence of CAR (**Fig. 2H-I**). Together, these results identify an early transitional stage of multi-potent T_H_ cells, which we term Th1/Tr1 progenitor (THR1p), that arises in secondary lymphoid organs during type 1 intestinal inflammation and that requires CAR to stabilize Tr1 cell lineage commitment.

### CAR promotes IL-27-mediated Tr1 cell development

Type 1 immunity involves naïve T_H_ cell priming by type 1 conventional dendritic cells (cDC1) that produce IL-12 and IL-27^32-34^. IL-12 signaling through the heterodimeric IL-12 receptor (IL-12R) evokes Th1 differentiation via Stat4, whereas IL-27-IL-27R signaling instigates differentiation of both Th1 and Tr1 cells via Stat1 and Stat3, respectively^35-37^. Given our finding that CAR-dependent ileal Tr1 cells developed predominantly during type 1 intestinal inflammation (**Fig. 1C-D**), we sought to examine interplay between CAR and IL-27 signaling during Tr1 cell development *in vitro*. CD45.1/CD45.2-mismatched wild type or CAR-deficient 10BiT reporter naïve T_H_ cells were purified from spleens of donor mice, mixed 1:1 and activated together (with crosslinking anti-CD3/anti-CD28 antibodies) in the same tissue culture wells with or without Th1-polarizing cytokines; expression of the 10BiT reporter was assessed on day 4, following two days of anti-CD3/anti-CD28 stimulation plus another two days of growth in IL-2-supplemented medium. As expected, IL-27 strongly amplified 10BiT reporter expression, compared with either IL-12 or media alone, and induction of 10BiT reporter expression by IL-27, but not IL-12, required CAR (**Fig. 3A-C**). Kinetic analyses revealed that naïve T_H_ cells activated in the presence of IL-27 upregulated CAR/*Nr1i3* gene expression within 24 hr; this was followed by CAR-dependent *Il10* upregulation that peaked at 48 hr post-activation and culminated with increasing cell surface Thy1.1 (10BiT) reporter expression on days 3-4 (**Fig. 3D-F**). Together, these results suggest that CAR contributes to initial IL-27-mediated priming of Tr1 differentiation, consistent with our *ex vivo* CITE-seq results that reveal peak CAR transcriptional activity in an early multi-potent Th1/Tr1 progenitor cell cluster (cluster 11 in **Fig. 2D-L**).

**Figure 3:**
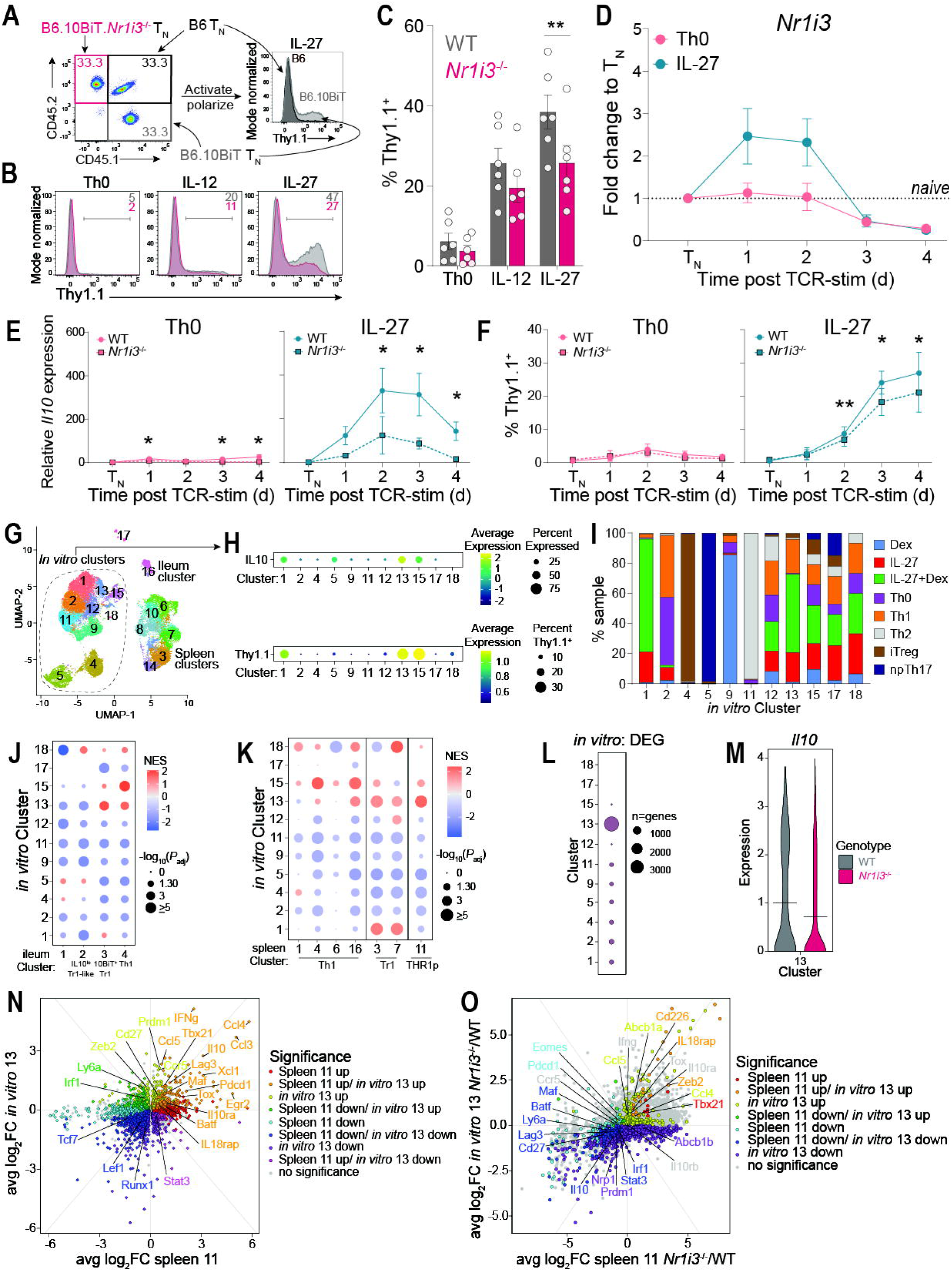
CAR promotes IL-27-induced Tr1 cell development. **(A-C)** *in vitro* activated and polarized CD4^+^ T cells in media alone (Th0), IL-12 (Th1), or IL-27. **(A)** Naïve CD4 T cells form 10BiT CAR-sufficient (WT, grey), CAR-deficient (*Nr1i3^-/-^*, pink) mice and non-10BiT WT B6 (B6, black) mice were mixed 1:1:1, activated and polarized. Cells were gated on viable, CD4^+^. **(B)** Representative histograms of Thy1.1 expression in WT (grey) and *Nr1i3^-/-^* (pink) cells. **(C)** Thy1.1 expression was measured. (Percent; n=6; +s.e.m.). \*\**P*<0.01; paired, one-way Anova with Greisser-Greenhouse correction. **(D)** Kinetic qPCR for *Nr1i3* in CAR-sufficient cells (fold change to WT naïve T cells (T_N_); n=4; ± s.e.m.). **(E)** Kinetic qPCR for *Il10* in CAR-sufficient (WT) and -deficient (*Nr1i3^-/-^*) cells (fold change to naïve T cells (T_N_); n=10; +s.e.m.). \**P*<0.05, \*\**P*<0.01, paired, parametric multiple t-tests. **(F)** Thy1.1 kinetics by flow cytometry in CAR-sufficient (WT) and -deficient (*Nr1i3*^-/-^) cells in Tr1 conditions. (Expression; n=6; + s.e.m.). \**P*<0.05, \*\**P*<0.01; paired, parametric multiple t-tests. **(G-O)** CITE-seq analysis of *in vitro* polarized WT and *Nr1i3^-/-^* cells. **(G)** UMAP projection of all hashed samples- *in vitro* polarized, ileum and spleen *ex vivo* isolated CD4^+^ T_H_ cells post adoptive Rag1^-/-^ transfer. **(H)** *Il10* and Thy1.1 expression in each cluster containing *in vitro* polarized cells. **(I)** Abundance of each CD4^+^ polarized T cell per *in vitro* cluster. **(J)** Geneset enrichment analysis (GSEA) between ileum subclusters and each *in vitro* cluster. **(K)** GSEA between spleen Th1, Tr1, and THR1p subclusters and *in vitro* clusters. **(L)** Differentially expressed genes (DEG) between WT and *Nr1i3^-/-^* cells in *in vitro* clusters. **(M)** Expression of *Il10* in *in vitro* shared Th1/Tr1 cluster (cluster 13) in WT and *Nr1i3^-/-^* cells. **(N-O)** Fold Change Fold Change plots of differentially expressed genes between CITE-seq samples **(N)** Spleen cluster 11 vs *in vitro* cluster 13, and **(O)** Spleen cluster 11 CARko/WT vs *in vitro* cluster 13 CARko/WT.

To explore the mechanisms and specificity of CAR function during IL-27-mediated Tr1 cell development *in vitro*, we expanded our CITE-seq studies to interrogate transcriptional features of co-cultured CAR-sufficient or CAR-deficient 10BiT reporter T_H_ cells stimulated *in vitro* for 48 hr under a variety of polarizing conditions. These conditions included IL-27 and were designed to evoke differentiation of canonical CD4^+^ effector and regulatory T cell lineages, including Tr1, Th1, Th2, Foxp3^+^ induced (i)Treg and Th17 cells (**Fig 3G-I, S2A-B**). Seurat clustering of all *in vitro*-polarized T_H_ cells discriminated 18 unique transcriptional states (**Fig. 3G**). Intriguingly, T_H_ cells stimulated with IL-27 segregated into three 10BiT^+^ clusters, including: (*i*) cluster 1, which displayed specific enrichment for genes expressed in CAR-dependent ileal Tr1 cells from mice with T cell transfer colitis; (*ii*) cluster 15, which showed transcriptional similarities with ileal Th1 cells from mice with T cell transfer colitis; and (*iii*) cluster 13, which expressed genes associated with both ileal Tr1- and Tr1-associated genes, as well as with the unique spleen THR1p cluster (**Fig. 3G-K, S2C**). As before with spleen THR1p (cluster 11) cells, *in vitro* IL-27-stimulated cluster 13 cells displayed a uniquely large number of differentially expressed genes between constituents with and without CAR; this included reduced expression of *Il10* and Tr1-associated TFs (*Stat3*, *Batf*, *Maf*, *Irf1*) in cells lacking CAR, and concomitant increases in the Th1-associated genes, *Tbx21* (T-bet) and *Il18rap* (**Fig. 3L-O**). Together, these results suggest that IL-27 plays a substantial role in generating the diverse array of Tr1 and Th1 cells produced during type 1 intestinal inflammation *in vivo*. This notably includes Th1/Tr1 progenitor (THR1p) cells in which CAR transcriptional activity is required to stabilize commitment towards the Tr1 lineage.

### Glucocorticoid receptor activation synergizes with IL-27 and CAR to augment Tr1 cell development

Glucocorticoid receptor (GR), like CAR, is a NR with clinically validated immunoregulatory functions^38^. GR is activated endogenously by cortisol, a glucocorticoid, and pharmacologically by corticosteroids, a widely prescribed class of medicines used to suppress autoimmune and chronic inflammatory disease flares^38^. Among its myriad immunoregulatory functions, GR has been shown to synergize with vitamin D receptor (VDR) in promoting Tr1 cell development^39^. In hepatocytes, GR *trans*-activates CAR/*Nr1i3* gene expression^40,41^. Given our results identifying a key early role for CAR in IL-27-induced Tr1 cell lineage commitment, we finally sought to understand how GR promotes Tr1 cell development via both CAR-dependent and CAR-independent mechanisms.

Using the same *in vitro* workflows as before, we found that dexamethasone (Dex)-mediated GR activation was sufficient to induce *in vitro* differentiation of naïve T_H_ cells into Foxp3^-^10BiT^+^ Tr1-like cells, albeit less than IL-27 alone, and that GR activation strongly synergized with IL-27 to amplify 10BiT reporter expression in Tr1-like cells (**Fig. 4A-B**). As expected, Dex-induced Tr1-like cell development required GR, but not Stat3, whereas the opposite was true for IL-27-mediated Tr1-like differentiation (**Fig. 4C**). CAR was required for induction of Il10 gene expression and expression of the 10BiT reporter in Tr1-like cells stimulated with Dex alone or IL-27 plus Dex (**Fig. 4D-E**). Consistent with the role of GR in hepatocyte CAR expression, Dex treatment augmented IL-27-dependent CAR/*Nri13* upregulation during T_H_ cell activation (**Fig. 4F**), suggesting at least part of GR’s capacity to promote Tr1-like cell development *in vitro* involves enhancement of CAR/*Nr1i3* gene expression.

**Figure 4:**
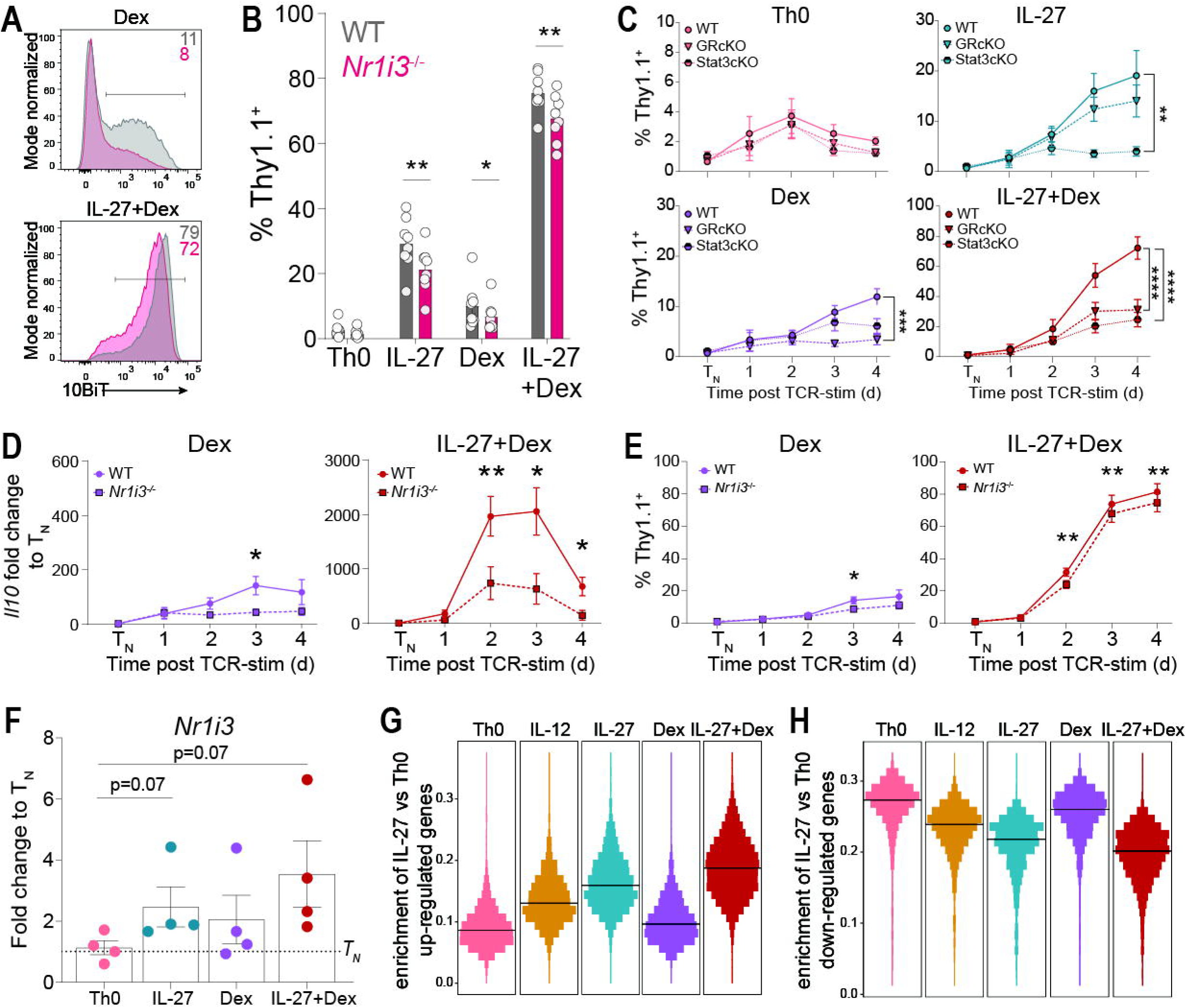
GR promotes CAR expression to augment IL-27-induced Tr1 cell development. **(A-C)** *in vitro* activated and polarized CD4^+^ T cells in media alone (Th0), IL-27, Dexamethasone (Dex), or IL-27+Dex. **(A)** Representative histograms of Thy1.1 expression in WT (grey) and *Nr1i3^-/-^* (pink) cells. **(B)** Thy1.1 expression was measured. (Percent; n=8; +s.e.m.). \**P<*0.05, \*\**P*<0.01; paired, one-way Anova with Greisser-Greenhouse correction. **(C)** Kinetic Thy1.1 expression by flow cytometry in CAR-sufficient (WT), GR^Flox/Flox^ distal LCKcre conditional CD4^+^ knockout (GRcKO), and Stat3^Flox/Flox^ distal LCKcre conditional CD4^+^ knockout (STAT3cKO) in Th0, IL-27, Dex and IL-27+Dex polarizing conditions. (Expression; n=5; ± s.e.m.). \*\**P*<0.01, \*\*\**P*<0.001, \*\*\*\**P*<0.0001; Mixed-effect model with Geisser-Greenhouse correction. **(D)** Kinetic qPCR for *Il10* in CAR-sufficient (WT) and -deficient (*Nr1i3^-/-^*) cells (fold change to naïve T cells (T_N_); n=10; +s.e.m.). \**P*<0.05, \*\**P*<0.01, paired, parametric multiple t-tests. **(E)** Thy1.1 kinetics by flow cytometry in CAR-sufficient (WT) and -deficient (*Nr1i3*^-/-^) cells in Tr1 conditions. (Expression; n=6; + s.e.m.). \**P*<0.05, \*\**P*<0.01; paired, parametric multiple t-tests. **(F)** Kinetic qPCR for *Nr1i3* 24-hours (Day 1, D1) post activation in CAR-sufficient cells (fold change to WT naïve T cells (T_N_); n=4; ± s.e.m.). **(G-H)** Enrichment of IL-27 mediated geneset, identified by comparing DEG of IL-27 to Th0 polarized cells compared to genes expresssed in Th0, IL-12, IL-27, Dex and IL-27+Dex polarized cells. **(G)** Enrichemt of IL-27 (vs Th0) mediated up-regulated genes. **(H)** Enrichment of IL-27 (vs Th0) mediated down-regulated genes.

Using CITE-seq to examine transcriptional profiles of Dex- and/or IL-27-stimulated T_H_ cells, we found that— aside from *Il10*—Dex and IL-27 alone produced few transcriptional similarities; Dex-treated T_H_ cells formed their own cluster (cluster 9) that separated on UMAP from the three IL-27-induced cell fates—Tr1-like (cluster 1), Th1-like (cluster 15) and THR1p-like (cluster 13) (**Fig. 3G-I**). At the same time, Dex-mediated GR activation strongly amplified IL-27-induced Tr1-like signatures, suggesting that IL-27 signaling (likely via Stat3) promotes large-scale chromatin accessibility upon which GR acts (**Fig. 4G-H)**. Accordingly, T_H_ cells activated in the presence of both IL-27 and Dex displayed representation within IL-27-associated Tr1-like (cluster 1) and THR1p-like (cluster 13) cells, but not Th1-like (cluster 15) cells (**Fig. 3G-I**). These results suggest that interplay between Stat3, GR and CAR shapes the balance of Tr1 *vs*. Th1 lineage commitment during type 1 intestinal inflammation. Contributions of GR may be particularly notable during patient responses to synthetic corticosteroids but may also be underappreciated during physiological type 1 intestinal inflammatory responses, given potential for both adrenal and extra-adrenal (intestinal) glucocorticoid production^24^.

### General conclusions

By mapping CAR-dependent T_H_ cell phenotypes evoked during discrete types of intestinal inflammation *in vivo*, and by unique T cell activation signals *in vitro*, we have shown that CAR acts in both early and lineage-biased manners to steer nascent Th1 cells towards the Tr1 lineage. CAR-mediated diversion of inflammatory Th1 cells into tolerogenic Tr1 cells likely evolved to balance the need for clearing enteric pathogens without inciting intestinal immune pathology. Indeed, the balance between Th1 and Tr1 cell appears particularly important in the ileum, where fewer Foxp3^+^ Treg cells are present compared with the colon^42^. Further, developmental relationships between Th1 and Tr1 cells during type 1 intestinal inflammation is akin to those shared by Treg and Th17 cells during type 3 immune responses. At a molecular level, our results suggest that much remains to be learned about the regulation and function of CAR, which remains viewed as a regulator of xenobiotic and endobiotic metabolism in the gastrointestinal tract. Specifically, if CAR activity in T_H_ cells followed the expected metabolism paradigm, one might assume—as we previously proposed—that CAR would act locally within the ileum lamina propria to enforce tolerogenic functions of mature Tr1 cells. Conversely, our results suggest that CAR expression and function in T_H_ cells peaks within the first day or two of antigen encounter and thus is most likely to act in secondary lymphoid structures, including the spleen, lymph nodes and Peyer’s patches. Toxic compounds known to activate CAR in gastrointestinal tissues should be less abundant in secondary lymphoid organs; thus, it is more likely that tolerogenic functions of CAR within developing Tr1 cells do not involve cytoplasmic sequestration or activation by currently understood xenobiotic/endobiotic ligands. Finally, our study suggests novel and actionable approaches to use CAR-modulating ligands to either promote or repress Tr1 cells during type 1 immune responses. In instances of Th1-related immune pathology, CAR agonist ligands could be leveraged to enhance abundance of developmentally related Tr1 cells. Conversely, CAR agonists could be pursued as a new approach to bolster Th1-assocaited immunity in settings of chronic viral infections or cancer.

## Methods

### Mice

All mice were C57BL/6J background and maintained under specific pathogen-free conditions at Scripps Florida and Dartmouth College. Importantly, there were no significant differences observed between the two institutions. Thy1.1-expressing IL-10 reporter, 10BiT^25^ mice, were provided by Casey Weaver (University of Alabama Birmingham) and bred with *Nr1i3^-/-^* mice^1^ provided by David Moore (University of California Berkley) and *Tnf^ΔARE/+^*mice^28^ provided by Fabio Cominelli (Case Western Reserve University). C57BL/6J CD45.1 (stock no: 002014), Rag1^-/-^ (stock no: 002216), distal LCK cre (stock no: 012837), GRflox (stock no: 012914), and Stat3flox (stock no: 016926) mice were purchased from The Jackson Laboratories. 10BiT *Nr1i3^-/-^ Tnf^ΔARE/+^* were generated by crossing 10BiT *Nr1i3^-/-^* mice to *Tnf^ΔARE/+^* mice. Sex matched mice were co-housed at weaning and analyzed at 20-weeks of age. All experiments and procedures were approved by the Institute Animal Care and Use Committees at Scripps Florida and Dartmouth College.

### CD4^+^ T cell isolation, activation and polarization

Purified CD4^+^CD25^-^ T cells were magnetically isolated from single cell suspensions prepared from spleen and peripheral lymph node mononuclear cells that were mechanically disrupted and passed through 10-µM nylon filters (BD Biosciences) using an EasySep magnetic T cell negative isolation kit (Stem Cell Technologies, Inc.; cat # 19852A) with the addition of a biotin anti-mouse CD25 antibody (0.5µg/mL; PC61; Biolegend).

Magnetically enriched CD4^+^ T cells were cultured in Dulbecco’s modified Eagle’s medium (DMEM) supplemented with 10% heat-inactivated fetal bovine serum (Gibco), 2mM L-glutamine (Gibco), 50 µM 2-mercaptoethanol (Sigma) 1% MEM vitamin solution (Gibco), 1% MEM non-essential amino acids solution (Gibco), 1% Sodium pyruvate (Gibco), 1% arg/asp/folic acid (Gibco), 1% HEPES (Gibco), 0.1% gentamicin (Gibco) and 100 U/mL Pen-Strep (Gibco).

Magnetically enriched CD4^+^CD25^-^ T cells were seeded (at 4x10^5^ cells/cm^2^ and 1x10^6^ cells/mL) in 96-well U- or flat-bottom plates pre-coated for 2hr at 37 °C with goat-anti hamster whole IgG (50µg/mL; Invitrogen). Activation was induced by adding hamster-anti-mouse CD3ε (0.3 or 1µg/mL; BioXcell) and hamster-anti-mouse CD28 (0.25 or 0.5 µg/mL; BioXcell). After 48 hr, cells were removed from coated wells and re-cultured at 1x10^6^ cells/mL in medium with or without 10 U/mL recombinant human IL-2 (rhIL-2) (NIH Biorepository), depending on the experiment. For polarization studies, cells were activated in the presence of the following sets of cytokines and/or neutralizing antibodies (all from R&D Systems unless specified): T_H_0: medium alone; T_H_1: recombinant huma (rh)IL-12 (5ng/mL) plus anti-mouse IL-4 (5µg/mL; Biolegend); T_H_2: rhIL-4 (10 ng/mL) plus anti-mouse IFNγ (5µg/mL; Biolegend); non-pathogenic (np) T_H_17: recombinant mouse (rm)IL-6 (40ng/mL) plust rhTGFβ1 (1ng/mL), anti-mouse IFNγ (5µg/mL; Biolegend) and anti-mouse IL-4 (5µg/mL; Biolegend); pathogenic (p)T_H_17: rmIL-6 (40ng/mL) plus rhTGFβ1 (1ng/mL), rhIL-23 (10ng/mL), anti-mouse IFNγ (5µg/mL; Biolegend) and anti-mouse IL-4 (5µg/mL; Biolegend); induced T regulatory (iT_reg_): rhTGFβ1 (5ng.mL) plus rhIL-2 (10U/mL), anti-mouse IFNγ (5µg/mL; Biolegend) and anti-mouse IL-4 (5µg/mL; Biolegend); IL-27: rhIL-27 (100ng/mL); Dex: Dexamethasone (100nM; Sigma-Aldrich); IL-27+Dex: rhIL-27 (100ng/mL) plus dexamethasone (100nM; Sigma-Aldrich). Cytokine, antibodies and/or dexamethasone were added at the time of activation (day 0) and re-added to expansion medium between days 2 and 4 of culture. Cells were analyzed for intracellular expression of transcription factors and/or cytokines, to confirm polarization, on day 2 or day 4 after re-stimulation with phorbol 12-myrisate 13-acetate (PMA; 10nM; Life Technologies) and ionomycin (1µM; Sigma-Aldrich) for 3-4 hr in the presence of brefeldin A (BFA; 10µg/mL; Life Technologies).

### Gut preps (Single cell suspensions of epithelial and lamina propria cells)

Preparation of single cell suspensions were performed as previously described^43^. Briefly, ileum (10cm) and colon were removed, opened longitudinally and rinsed with ice-cold PBS. Peyer’s patches were removed from the ileum. Dissected ileum and colon were cut into pieces (∼1-2cm) and epithelial cells were dissociated by incubating in RPMI medium containing 1% FBS, 1 M MgCl_2_ and 0.5 M EDTA with shaking at 280 rpm for 20 min at 37 °C, twice. The remaining tissues were digested in RPMI medium containing 20% FBS and 0.2 mg/mL Collagenase III (ThermoFisher Scientific) with shaking at 280 rpm for 37 min at 37 °C, twice. The epithelial and lamina propria cell fractions were filtered through 100-µM cell strainers. Lamina propria cells were resuspended in 40% Percoll solution with centrifugation at 3,000 g for 12 min at room temperature. Cells were then washed and resuspended in PBS containing 2% FBS.

### T cell transfer colitis

For mixed congenic T cell transfers, magnetically isolated T cells (CD4^+^CD25^-^) from B6-derived 10BiT wild-type and 10BiT *Nr1i3*-deficient T cells were mixed 1:1 and transferred together (0.5x10^6^ total cells) intraperitoneally (i.p.) into syngeneic *Rag1^-/-^*recipients and analyzed 2.5 weeks after transfer. All *Rag1^-/-^*recipients were weighed immediately before T cell transfer to determine baseline weight and then weighed weekly after T cell transfer for the duration of the experiment. Transferred *Rag1^-/-^* mice were killed when they had lost 20% of pre-transfer baseline weight. All *Rag1^-/-^*mice that received different donor T cells were co-housed to normalize microflora exposure.

### Flow cytometry

Cell surface and intracellular FACS stains were performed at 4 °C for 20 min, washed with phosphate buffered saline (PBS) and acquired on a flow cytometer. Dead cells were excluded with eFluor 780 Viability stain. Anti-mouse antibodies used for FACS analysis included: APC anti-CD45.1 (A20), PE anti-CD4 (RM4-5), BV510 anti-CD4 (RM4-5), BUV395 anti-CD4 (RM4-5), BV510 anti-CD25 (PC61), PE anti-CD25 (PC61), BV650 anti-CD44 (IM7), BUV496 anti-TCRβ (H57-597), FITC anti-CD45.2 (104), Percp-Cy5.5 anti-Thy1.1 (OX-7), BV605 anti-TNFα (MP6-XT22), PE-Cy7 anti-IL-10 (JES5-16E3), BV711 anti-IFNγ (XMG1.2), APC anti-IFNγ (XMG1.2), BV421 anti-IFNγ (XMG1.2), PE anti-IL-4 (11B11), Alexa Flour700 anti-IL-17a (TC11-18H10.1), FITC anti-IL-17a (TC11-18H10.1), PE-CF594 anti-RORγt (Q31-378), eFluor 450 anti-FoxP3 (FJK-16s), and BV421 anti-IL-2 (JES6-5H4). Cells were analyzed for intracellular expression of Transcription factor staining buffer set following manufacturer’s instructions (ThermoFisher Scientific) or homemade cytokine staining kit, to confirm polarization, on day 2 or day 4 after re-stimulation with phorbol 12-myrisate 13-acetate (PMA; 10nM; Life Technologies) and ionomycin (1µM; Sigma-Aldrich) for 3-4 hr in the presence of brefeldin A (BFA; 10µg/mL; Life Technologies).

### qPCR

RNA was isolated from cultured cells using RNeasy Mini columns with on-column DNase treatment (Qiagen). RNA was used to synthesize cDNA with a high capacity cDNA reverse transcription kit (Life Technologies). Taqman qPCR was performed on a StepOnePlus real time PCR instrument (Life Technologies/Applied Biosystems) or QuantStudio 6 pro (ThermoFisher Scientific) using commercial Taqman primer/probe sets (Life Technologies). Probes for mouse genes included: *Nr1i3* (Mm01283980_g1), *IL10* (Mm01288386_m1), and *ACTB* (Mm00607939_s1).

### Cellular Indexing of Transcriptomes and Epitopes by sequencing (CITE-seq)

#### Single-cell suspension and library preparation

Cellular Indexing of Transcriptomes and Epitopes by sequencing (CITE-seq) was performed on magnetically enriched CD4^+^ T cells from spleen and ileum lamina propria (as described above) cells of *Rag1^-/-^* mice injected 2.5 weeks earlier with 1:1 mixtures of 10BiT CD45.1 wild-type and 10BiT CD45.2 *Nr1i3^-/-^* naïve T cells, as well as cultured T cells following 48 hours activation (T_H_0, T_H_1, T_H_2, npT_H_17, iTreg, IL-27, Dex, and T_R_1). Preparation of single cell suspensions were performed as previously described in CITE-seq and Cell hashing protocol from NY Genome Center Technology Lab (https://citeseq.wordpress.com/wp-content/uploads/2019/02/cite-seq_and_hashing_protocol_190213.pdf). Briefly, 1-2x10^6^ cells from each sample were suspended in staining buffer (2%BSA/ 0.01% Tween, PBS; Biolegend) with the addition of 10uL of FC Blocking reagent (FCX; Biolegend) for 10 min at 4 °C. Totalseq A cell hashtag (HTO) and Antibody derived tag (ADT) antibodies were used (Biolegend). 2x ADT pool was prepared for 10 samples containing CD45.1 (A20), CD45.2 (104), and CD90.1 (Thy1.1, OX-7). 0.5µg of unique cell hashing antibodies were added to each sample (Totalseq-A0301-A0310; M1/42, 30-F11) for 30 min at 4 °C. Cells were washed 3 times, filtered with 40-µM nylon filter, counted and viability checked for ≥ 80%. Samples were mixed at equal cell numbers for library preparation. Libraries of the same cell mixes were prepared for 2 lanes of super-loaded 35,000 cells/lane using 10x kit. Separate ADT and HTO-derived cDNA and mRNA-derived cDNA were amplified, purified, and sequenced.

### Preprocessing and Demultiplexing

Hashed single cell RNAseq data was collected in two wells. Analysis was performed separately for each of the two wells, until being merged later in the methods. 10X sequencing data was loaded into Seurat 5.3.0^44^. The sequencing data was split into three assays of features: RNAseq features containing RNA-sequencing reads, HTO features containing cite-seq markers to determine cell type, and ADT features containing cite-seq markers to determine genotype and Thy1.1 expression. CITE-seq data was collected and normalized using centered log-ratio transformation. Cells were demultiplexed using Seurat MULTIseqDemux^44^, using an automatically determined threshold, demultiplexing cell type (HTO) then genotype (ADT). The following cell types were multiplexed by HTO and mapped to HT01=HT)10: Th0, Th1, Th2, npTh17, iTreg, IL-27, Dex, IL-27+Dex, *ex vivo* spleen, and *ex vivo* ileum. WT cells were mapped to ADT CD45.1 and *Nr1i3^-/-^* cells were mapped to CD45.2. An additional ADT, CD90.1, was used to measure the IL-10 Thy1.1-expressing 10BiT reporter expression. Cells determined by MULTIseqDemux to be either doublets or negative in the HTO demultiplexing step were removed. The two wells were then merged. Quality control metrics for RNAseq data was calculated. Cells with counts between 800 and 15000, and with features between 500 and 5000, and whose counts were less than 5% mitochondrial were retained, while the others were removed. Genes which had at least one count in at least 10 cells were retained, while the others were removed.

### Normalization and Clustering

Cells were log-normalized using Seurat NormalizeData^44^, using a scaling factor of 10,000, then scaled using Seurat ScaleData^44^. Cell cycle was determined using Seurat CellCycleScoring^44^. Mouse homologs for Human S and g2m features present in Seurat’s human s and g2m references contained in Seurat ‘cc.genes.updated.2019’^44^ which reference Ensembl V105^45^. The top 5000 highly-variable genes were detected with Seurat FindVariableFeatures^44^ using variance-stabilizing transformation. Counts were scaled and the top 50 PCs were calculated using Seurat ScaleData and RunPCA^44^. Cell cycle was estimated using the cell-cycle genes specified in Seurat cc.genes.updated.2019^44,46^. Conversions between human and mouse genes used biomaRt getLDS^47^ and Ensembl V105^45^. Gene counts were again normalized using log-normalization, and scaled using Seurat::SCTransform^44^ using the “v2” variance-stabilizing transformation. The top 5000 variable features were calculated and the effect of counts, features and cell cycle were regressed from the dataset. The first 50 principal components were calculated, and a SNN graph was calculated from it using Seurat::FindNeighbors^44^. A final UMAP was calculated using Seurat RunUMAP^44,48^ based on the SNN graph. Leiden clustering was performed using Seurat FindClusters^44,49^, and optimum resolution was determined to be 0.6, evaluated using clustree^50^ to determine the stability of the clusters at each resolution. Cell genotypes were demultiplexed using ADT again, and this time cells determined by MULTIseqDemux to be doublets or negative for the ADT assay were removed. After clustering had been performed, the normalized counts using Seurat SCTransform^44^ were discarded.

### Differential Gene Expression and Gene Set Enrichment

Data was centered and scaled again using Seurat ScaleData^44^. Highly variable genes re-calculated using Seurat FindVariableFeatures^44^ to select 5000 variable genes using variance stabilizing transformation. Cluster markers were calculated using the Wilcox test on the new normalized counts, using Seurat FindAllMarkers^44^. Marker genes between WT and CARko cells were calculated for each cluster using wilcox tests and SeuratFindMarkers^44^, performed separately for each cell-type and leiden cluster separately. p-values were normalized using q-values. Gene sets were taken from the “Biological Process” gene-sets of the Gene Ontology go-basic database^51,52^ (https://zenodo.org/records/18422732). The database was filtered to limit the gene sets to those that contained between 20 and 2000 genes. Additional gene sets were loaded from MSigDB v2024.1 mouse database^53,54^ and from IL-10 expressing T cell subset publications^14,19,20,27,29,55^ to help identify cell-types. The custom gene sets were appended to the go-basic database^51,52^ (https://zenodo.org/records/18422732). An additional set of gene sets was loaded from reactome.db^56^, using gene sets containing 10 and 500 genes. Gene Set Enrichment (GSE) was calculated using fgsea method with clusterProfiler GSEA^57,58^.

### Ileum and Spleen Subclustering, DGE, and GSE

The ileum and spleen samples were separated from the rest of the datasets to analyze individually. Normalization, scaling, highly variable genes, shared-nearest-neighbor, and leiden clustering were all re-performed on the ileum and spleen individually, separate from the previous analysis. Sub-clustering was performed using SCTransform flavor “v2”^44^ and selecting the top 5000 variable features. Read counts and phase were regressed out of the ileum sample, while phase was regressed out of the spleen sample with the Harmony package^59^. The top 20 and 30 PCs were selected for ileum and spleen samples respectively. Leiden clustering was performed^49^, and clustree analysis^50^ found that the leiden resolution of 1 for ileum and 1.5 for spleen to produce stable clusters. Marker gene detection and gene set enrichment were calculated again using the ileum and spleen subclusters. Marker genes from ileum and spleen subclusters were saved and used as gene sets for additional GSE analysis. For plots comparing two sets of fold-changes, for each group being compared, the fold-change and significance based on q-values of the differential expression for inside the group was compared to cells outside the group. Cells were colored based on their significance across both groups.

### Velocity analysis

Data was converted from Seurat^44^ to Anndata^60^ using SeuratDisk (https://mojaveazure.github.io/seurat-disk/reference/SeuratDisk-package.html). RNA splicing was calculated with Velocyto^61^ using the GENCODE vM23 reference genome^62^, and masking regions from the UCSC mm10 repeat masker dataset^63^. RNA velocity was calculated with scvelo using “dynamical” modeling^64^. Clusters relationships were graphed using PAGA^65^, and velocity analysis from scvelo was used to assign directionality to the graph^64^. Connections between clusters in pAGA graphs were filtered to display connection strengths above 0.1.

### Statistical analyses

Statistical analyses for were performed using Prism (GraphPad) for Figures 1-2 and *R* for Figure 3. *P*values were determined by paired or unpaired Student’s t-tests, one-way ANOVA, Wilcoxon Rank Sum test, or mixed effect model (two-way ANOVA) as appropriate and as listed in the figure legends. The statistical significances of differences (\**P*<0.05, \*\**P*<0.01, \*\*\**P*<0.001, \*\*\*\**P*<0.0001) are specified through figures and legends. Unless otherwise noted in legends, data are shown as mean ± s.e.m. P-value adjustment for CITE-seq for multiple testing was performed using Q-value adjustment^66^.

## Abbreviations

CAR: constitutive androstane receptor (*Nr1i3*)
Th1: CD4 T helper 1 cells
Tr1: Type 1 regulatory T cell
Th17: CD4 T helper 17 cells
NR: nuclear receptor
SI: small intestine
T_H_: CD4+ T helper cell
IL-10: interleukin 10
BA: bile acids
XRE: xenobiotic response element
IL-27: interleukin 27
Dex: Dexamethasone
IL-12: interleukin 12
LBD: Ligand Binding Domain
FoxP3: Forkhead box 3
Stat3: Signal transducer and activator of transcription 3
GR: Glucocorticoid receptor (*Nr3c1*)

## Figure legends

**Supplemental Figure 1: CAR-dependent T cell phenotypes in T cell transfer colitis and TNFΔARE ileitis. (A-G)** *in vivo* transfer of CAR-sufficient (WT) and CAR-deficient (*Nr1i3^-/-^*) 10BiT naïve CD4+ T cells into Rag1-deficient (Rag1^-/-^) hosts. Cells were isolated from the colon and spleen 2.5 weeks post transfer, stained and analyzed via flow cytometry. **(A)** Cells were gated on CD4^+^ T_H_ cells, CD44^+^ FoxP3^-^ and gated on congenic marker, CD45.1^+^ WT and CD45.1^-^ *Nr1i3^-/-^*. **(B)** Representative flow cytometry plot Thy1.1 vs FoxP3, gated on CD44^+^ cells. **(C)** Expression of Thy1.1 was measured. (Percent; n=11; ± s.e.m.) \*\*\**P<*0.001, paired, parametric two-tailed Student’s *t-*test. **(D)** Expression of IL-10 was measured. (Percent; n=11; ± s.e.m.) \*\*\**P<*0.001, paired, parametric two-tailed Student’s *t-*test. **(E)** Representative flow cytometry plot IFNγ vs IL-17a. **(F)** Expression of IFNγ was measured. (Percent; n=11; ± s.e.m.) \**P<*0.05, paired, parametric two-tailed Student’s *t-*test. **(G)** Expression of IL-17a was measured. (Percent; n=11; ± s.e.m.) \**P<*0.05, paired, parametric two-tailed Student’s *t-*test. **(H-N)** Spontaneous ileitis TNF^ΔARE/+^ mice crossed to 10BiT CAR-sufficient and CAR-deficient mice were cohoused until 20 weeks of age. Cells were isolated from the colon and spleen, stained and analyzed via flow cytometry. **(H)** Cells were gated on viable, CD4^+^ T_H_ cells, CD44^+^ FoxP3^-^. **(I)** Representative flow cytometry plot Thy1.1 vs FoxP3, gated on CD44^+^ cells. **(J)** Expression of Thy1.1 was measured. (Percent; n=8; +s.e.m.), paired, one-way Anova with Greisser-Greenhouse correction. **(K)** Expression of IL-10 was measured. (Percent; n=8; +s.e.m.), paired, one-way Anova with Greisser-Greenhouse correction. **(L)** Representative flow cytometry plot IFNγ vs IL-17a. **(M)** Expression of IFNγ was measured. (Percent; n=8; +s.e.m.), paired, one-way Anova with Greisser-Greenhouse correction. **(N)** Expression of IL-17a was measured. (Percent; n=8; +s.e.m.), paired, one-way Anova with Greisser-Greenhouse correction.

**Supplemental Figure 2: Cytokine and gene expression in *in vitro*-polarized T_H_ cell subsets. (a-b)** Naïve CD4^+^ T cells isolated from triple reporter FoxP3^RFP^ IL-17a^GFP^ 10BiT^+^ IL-10 reporter mice were polarized into Th0, Th1, Th2, npTh17, pTh17, iTreg, and IL-27+Dex Tr1 conditions for 4 days. Cells were then stimulated and analyzed via flow. **(a)** Representative flow plots of intracellular markers and surface reporters. **(b)** Percent of intracellular markers and surface reporters quantified. (Expression; n=4, ± s.e.m.). **(c)** Expression of key Tr1- and Th1-associated genes in clusters containing cells polarized *in vitro*.

## Notes

### Competing Interest Statement

The authors have declared no competing interest.

### Funding

- National Institute of Allergy and Infectious Diseases: R01AI143821, R01AI164772, U01AI163063

