## Supplementary figures and images for "Constitutive androstane receptor directs developing Th1 cells towards the Tr1 lineage"

### Supplemental Figure 1

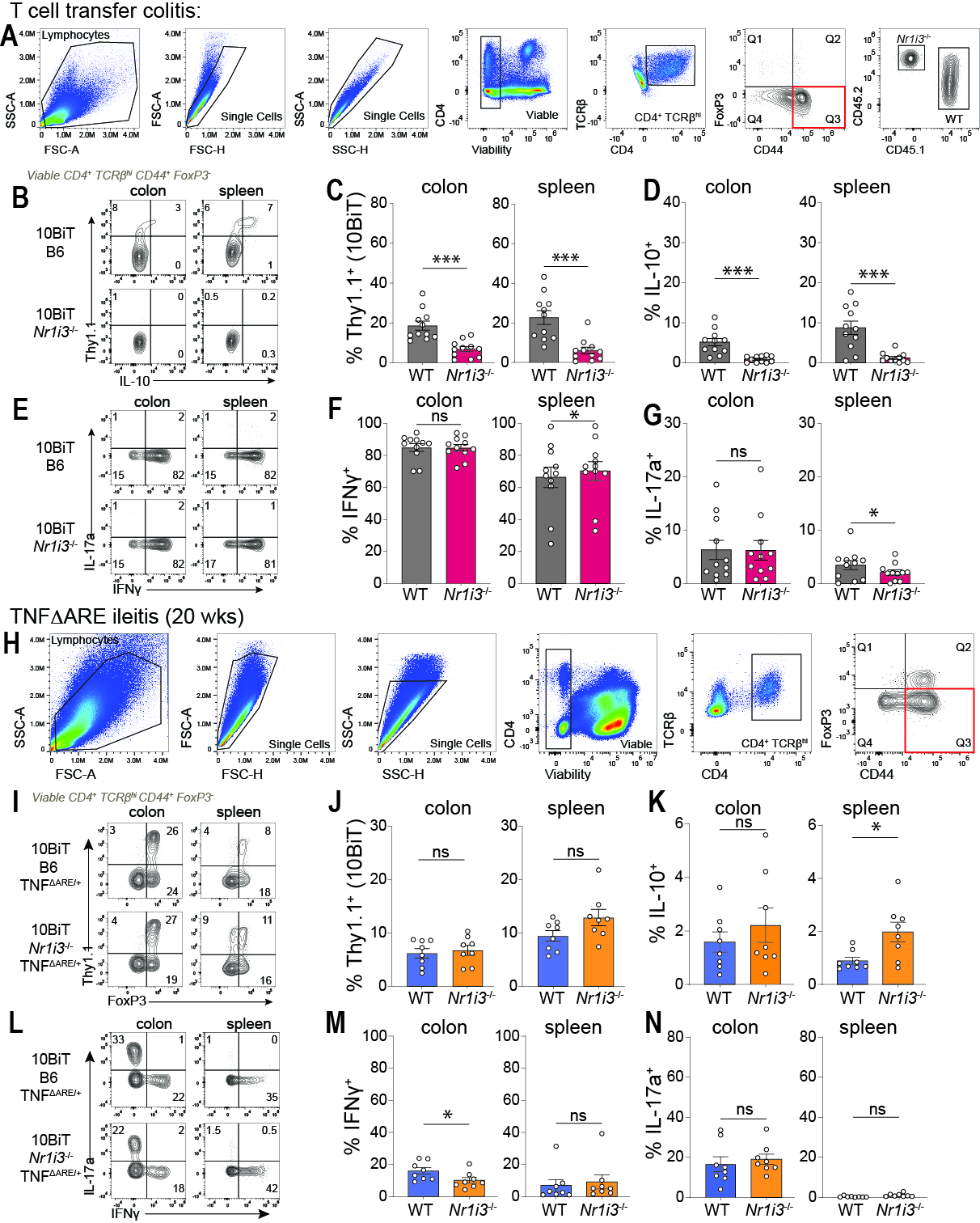

### Supplemental Figure 2

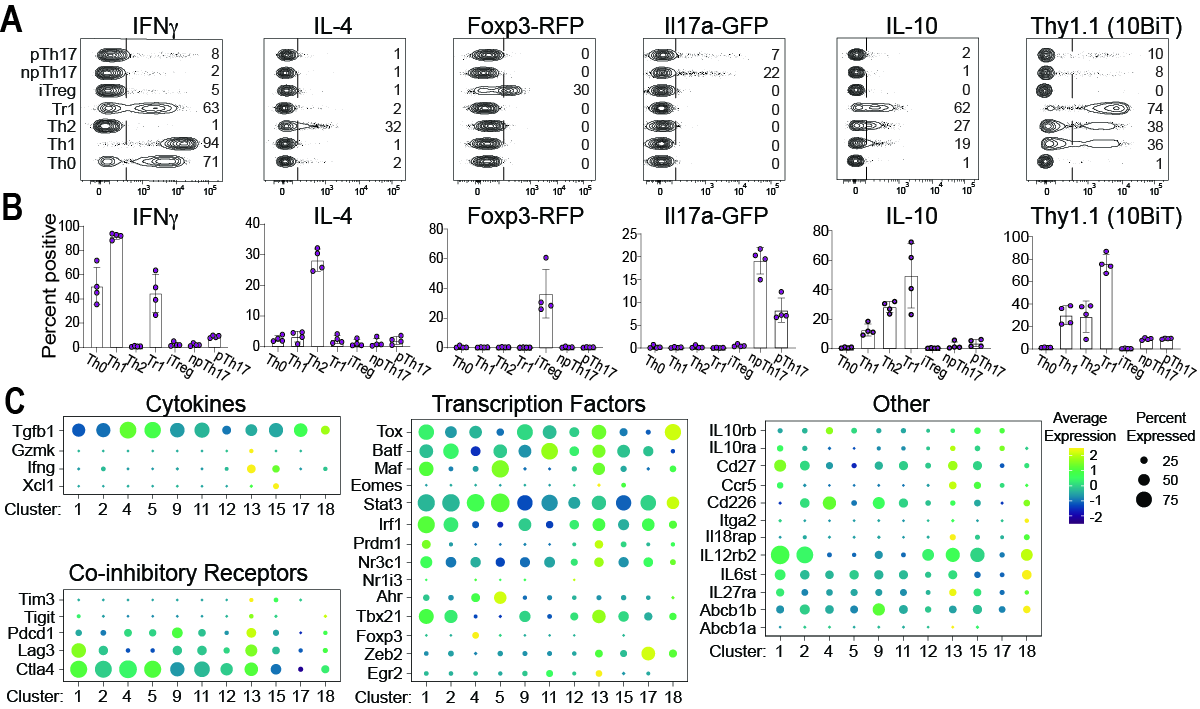
